# RNA-Ribo Explorer: interactive mining and visualisation of Ribosome profiling data

**DOI:** 10.1101/2021.03.23.436679

**Authors:** D. Paulet, A. David, E. Rivals

## Abstract

RNA translation has long been thought as a stable and uniform process by which a ribosome produces a protein encoded by the main Open Reading Frame (ORF) of an mRNA. Recently, growing evidence support incomplete correlation between RNA and protein abundance levels, the existence of alternative ORFs in numerous mammalian RNAs, and the involvement of ribosomes in gene expression regulation, thereby challenging previous views of translation. Ribosome profiling (aka Ribo-seq) has renewed the study of translation by enabling the mapping of translating ribosomes on the whole transcriptome using deep-sequencing.

Despite increasing use of Ribo-seq, recent review articles conclude that flexible, interactive tools for mining such data are missing. As Ribo-seq protocols still evolve, flexibility is highly desirable for the end-user. Here we describe RNA-Ribo-Explorer (RRE) a stand-alone tool that fills this gap. With RRE, one can explore read-count profiles of RNAs obtained after mapping, compare them between conditions, and visualize the profiles of individual RNAs. Importantly, the user can mine the data by defining queries that combine several criteria to detect interesting subsets of RNAs. For instance, one can ask RRE to find all RNAs whose translation of UTR region compared to that of the main ORF has changed between two conditions. This feature seems useful for finding candidate RNAs whose translation status or processing has changed across conditions.

RRE is a platform independent software and is freely available at https://gite.lirmm.fr/rivals/RRE/-/releases.

## 1 Introduction

Translation regulation refers to alterations of ribosome behavior that influence the process of translation and thus impacts the final protein level of a gene. Translation is an intricate process performed by ribosomal complexes. In eukaryotes, the ribosome scans the RNA from the cap, finds a start codon which triggers translation until the end of an Open Reading Frame (ORF) [18]. This scanning model has been challenged by the existence of ORFs located upstream or downstream the main ORF (in regions described as UnsTranslated Regions or UTRs for short). Most of these alternative ORFs in eukaryotes are located between the cap and the main ORF start codon (i.e., in the 5’UTR). Now thousands of such alternative ORFs located in UTRs have been found in mammalian transcriptomes [13], and mutations that create or alter such ORFs were proven to control the protein synthesis of the main ORF [18]. The well studied case of ATF4 mRNA in humans shows that several upstream ORFs are exquisitely combined to achieve a precise regulation of that gene [20]. Mutations in untranslated region can create, shorten or lengthen upstream ORFs, and induce dysregulation of the main ORF, which may in turn cause pathological disorders, for instance, in hereditary melanoma [7]. Actually, when in response to a stress, a cell turns down translation of most genes, but increases the peptide synthesis of a subset of “response” genes, it often controls the latter by bypassing upstream ORF [19]. All these phenomena emphasize the importance of translational regulation of gene expression. However, the precise mechanisms behind translational regulation require further investigation. The advent of *Ribosome profiling* or *Ribo-seq* has opened new avenues in this direction. A fine understanding of translational regulation may also help improving gene expression for applications in synthetic biology [11].

### 1.1 Ribo-seq

RNA-seq allows to explore the transcriptome – both qualitatively and quantitatively – and to uncover its astonishing variability, while quantitative proteomics can provide estimates of protein abundance at large scale. However, it is still challenging to predict protein abundance from transcript quantities, for the intermediate step of translation participates in regulating gene expression more deeply than previously thought [5]. Ribosome profiling, also termed Ribosome sequencing or Ribo-seq, is a recent Next Generation Sequencing (NGS) assay that enables us to interrogate the translation process in a transcriptome wide manner.

Ribo-seq allows to map the positions of translating ribosomes on the entire transcriptomes. Current protocols follow multiple steps, among which are i/ a cell treatment with a drug that inhibits either elongating or initiating ribosomes, ii/ application of nuclease to digest RNA portions not covered by ribosomes and capture of Ribosome Protected Fragments (RPF), iii/ fragment selection on size, rRNA depletion, conversion into DNA, library preparation and deep sequencing [4].

Despite the fact that Ribo-seq remains a challenging technique and suffers from some biases, it became the main molecular assay to study translation at large scale and is increasingly used to answer a variety of questions. Notable aspects of translation studied until now include i/ the quantification of translational control, ii/ the identification of alternative Open Reading Frames (short ORFs, upstream ORF located in 5’UTR, etc) in peptidomics or functional genomics for instance, iii/ the dynamics and mechanics of translation [2]. Research based on Ribo-seq uncovered that translation is more pervasive in mammals than previously thought [4]. Existing Ribo-seq datasets show that Ribo-seq profiles differ from RNA-seq profiles, because first read accumulation strongly depends on the kinetic of translation, and read coverage is generally lower than in RNA-seq (as many reads match ribosomal RNA sequences). Moreover, Ribo-seq reads are short (usually 25-32 nucleotide long), exhibit a trinucleotidic periodicity pattern related to the coding frame, and require filtering of reads coming from ribosomal RNAs. Due to its specificities, Ribo-seq require adadpted analysis software.

### 1.2 Tools for Ribo-seq analysis

A wide range of computational methods have been developed for analyzing ribosome profiling datasets:

- to pipeline the primary analysis Ribo-seq reads, from mapping to read counts per transcripts [10],
- to collate or view footprint profiles for known RNAs [12], to normalize the read counts [14], or to estimate translation efficiency relatively to transcript abundance [6],
- to postprocess Ribo-seq profiles to predict distinct features like alternative or small ORFs [1, 10], ribosome stalling events, or estimate Codon Usage Bias [16].

A nice survey on methods for predicting ORFs was published in October 2017 [2], while a more comprehensive overview of computational resources for all kinds of analyses appeared in January 2019 [21], witnessing a booming interest in ribosome profiling. Available computational methods for Ribo-seq data often concentrate on one precise aspect (like removing of contaminant reads [9], or estimation of the position offset [3]), are implemented in different frameworks (R package, standalone programs, or Galaxy tools), which does not facilitate their combination and inter-operability. Noteworthy, both of the above-mentioned review articles underline the lack of tool for mining and exploring Ribo-seq data interactively. As the protocols of Ribo-seq assays are still evolving, their yield, reproducibility, and biases hinder the establishment of standards in computational analysis of Ribo-seq. The diversity of primary read analysis pipelines reflect the need to leverage Ribo-seq in different applications.

We therefore reason that it is crucial for the end user to be able to visualize and mine its Ribo-seq datasets in a simple, flexible and interactive way. Despite the availability of dedicated read processing pipelines or of visualization solutions based on genome viewers [2, 21], a tool for this task is still missing. Hence, we propose RNA-Ribo Explorer, a stand alone software for visualization and mining of Ribo-seq datasets in transcriptome wide studies.

## 2 Material

To demonstrate the functionalities of RNA-Ribo Explorer, we use two publicly available Ribo-seq datasets: 1) one compares human tumoral vs normal cells from a fresh kidney carcinoma [8], 2) the other contrasts human embryonic kidney (HEK) cell lines expressing DDX3 mutant or not in normal and arsenite-induced stress conditions [15]. For simplicity, we will refer to the former as the *carcinoma* dataset, and to the later as the *HEK* dataset. Both data comes from Gene Expression Omnibus repository with accession number GSE70804 and GSE59821, respectively.

## 3 Method

### 3.1 Overview

RRE is an interactive and flexible tool that enables the user to explore and analyze ribosome profiling data in a dynamic, visual, and easy manner. RRE is intended for the biologist for it allows her/him to design queries exactly matching his/her needs, to obtain the query results as selection or plots, to export these in files, and to examine graphically the profiles for any desired mRNA. The user can also set key parameters interactively, which will refine the results on-the-fly, giving her/him the possibility to test hypotheses dynamically. Furthermore, RRE requires little know-how in computational biology, has a simple installation procedure, and is platform independent (for it is programmed in Java language). RRE provides

- specific features required for analyzing Ribo-seq,
- interactive visualization of Ribo-seq profile for any mRNA,
- data mining capabilities which the user performs by running three major query types.

RRE enables interactive data mining of RNA-seq and Ribo-seq, once the raw reads have been processed using a map-and-count pipeline. RRE takes as input specific files containing read counts, which can be generated from a BAM file using our script (Figure 1).

**Figure 1:**
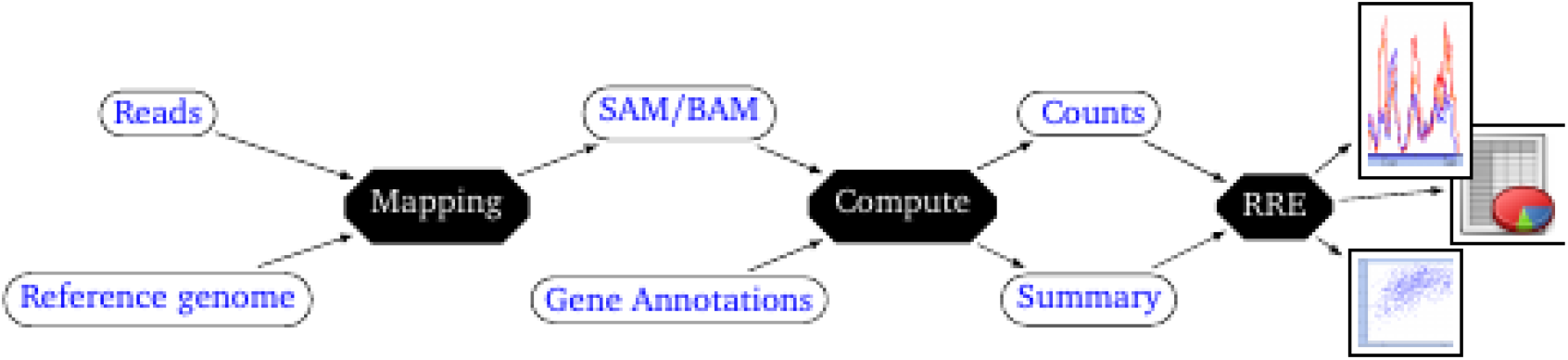
Overview of the read processing pipeline and position of RRE as a secondary analysis tool. Raw reads are mapped on the reference genome (or transcriptome), which yields a SAM/BAM file. This file combined with gene/RNA annotations allows computing read coverage counts for each position of each RNA, and separate counts in region annotated as untranslated or translated. This generates a count file, and a summary file that contains a subset of useful transcriptome annotations: these two files are light compared to the original BAM file and GCF files, and serve as input to RRE. RRE is then used for interactive data mining and visualisation in downstream analyses by the end user.

### 3.2 Ribo-seq specific features

In Ribo-seq, reads represent RPFs whose expected length is 25-32 nucleotides. As a RPF denotes the presence of a ribosome interacting with an mRNA, the distribution of reads mapped on a transcript should accumulate at the start codons and also exhibit a trinucleotidic periodicity. RRE provides functionalities to perform classical quality controls on each Ribo-seq dataset. The user can plot for instance the distribution of mapped read lengths to check whether they respect the expected distribution (see Figure 5 in documentation), or generate a metagene plot of read coverage per position relative to the start position of the main ORF (Figure 2) in order to verify the existence of a peak near the start codon and of the trinucleotidic periodicity pattern. To derive the position of the ribosome P-site (and hence of the decoded codon) from the mapping position of a read, one needs to add an offset, which is often set to 12 nucleotides in most studies. However, this offset may depend on the data set, on the protocol, and especially of the ribonuclease used [2]. Using the coverage values of all mRNAs for a given dataset, RRE will propose a suitable offset to the user and update the metagene read coverage plot taking into account the chosen offset value. The user can try several values and validate the most appropriate one, checking whether or not it improves e.g. the trinucleotidic periodicity pattern.

**Figure 2:**
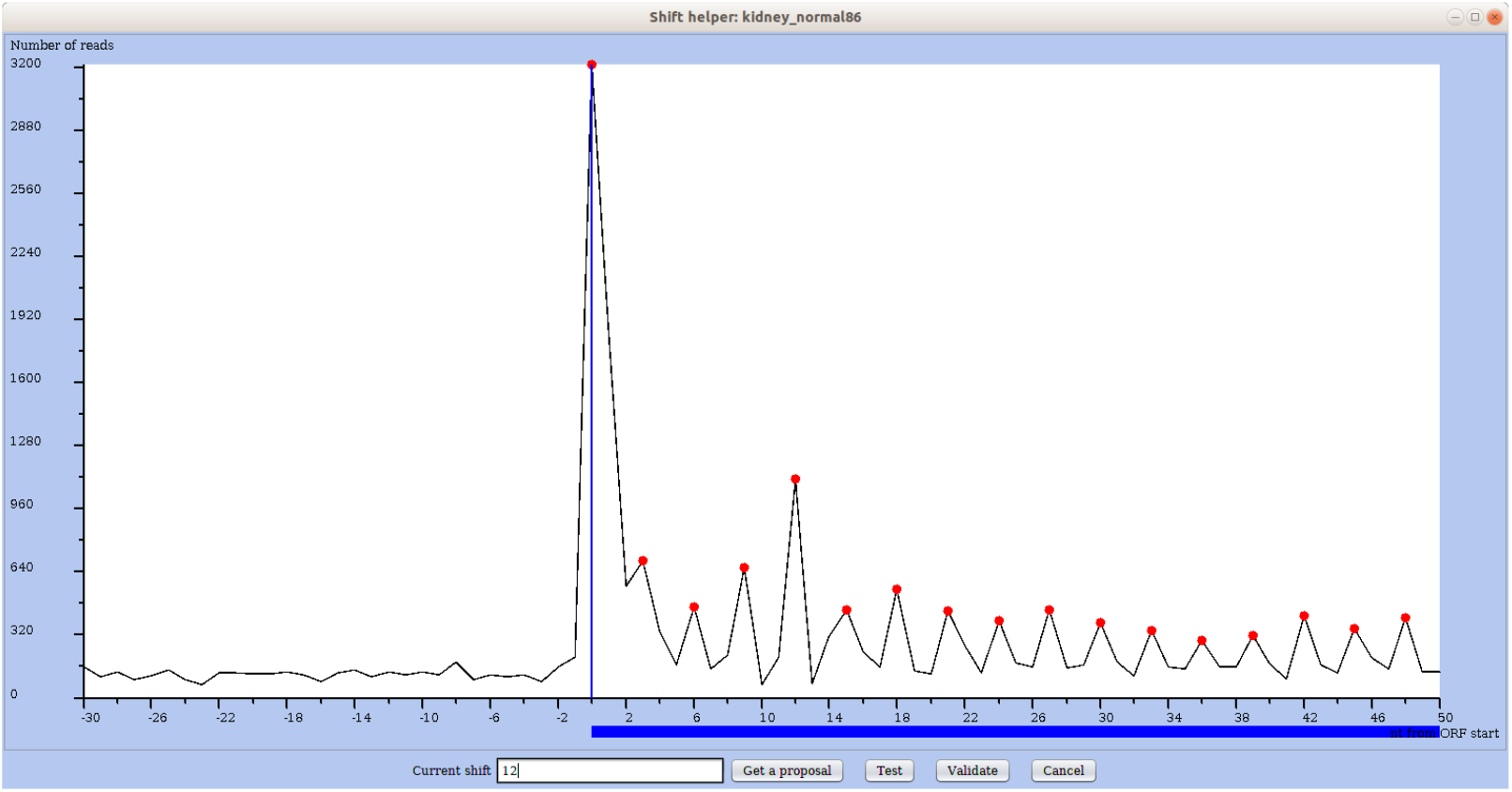
Metagene plot of read coverage over all mRNAs for the region around the start of the main ORF. RNA positions on the x-axis are relative to the start of the annotated main ORF (position 0). The red dots mark each third position from the start of the main ORF (0, 3, 6, etc). In the footer, the user can ask RRE to propose an offset/shift value specifically for this dataset or he/she can provide a value. The offset value is used to determine the P-site from the mapping position of a read (see User’s manual).

Note that such quality control and preprocessing steps can each be performed using some existing packages or software, while RRE integrates these functionalities in a single tool for the convenience of end users.

### 3.3 Interactive visualization of an mRNA profile

Figure 3 shows the profile window for the mRNA named P4HB using four selected conditions. The plot displays as many profiles for this mRNA as selected conditions and indicates the reference CDS as a blue bar (below the plot), the 5’ and 3’ UTRs being to the left and right of it. Once the user has selected one mRNA (using a search query or the result of a mining query), RRE lets him on-the-fly act on plot parameters, select or deselect conditions, select the range of read lengths considered in the plot, and view the mRNA’s rank in the distribution of coverages across all mRNAs. With such a multi-profile plot, the eye easily detects whether peaks and valleys have conserved positions across conditions (as in Figure 3) or not (as in Figure 8). RRE offers the possibility of zooming and moving along the sequence position (with the control panel in lower left corner of the window).

**Figure 3:**
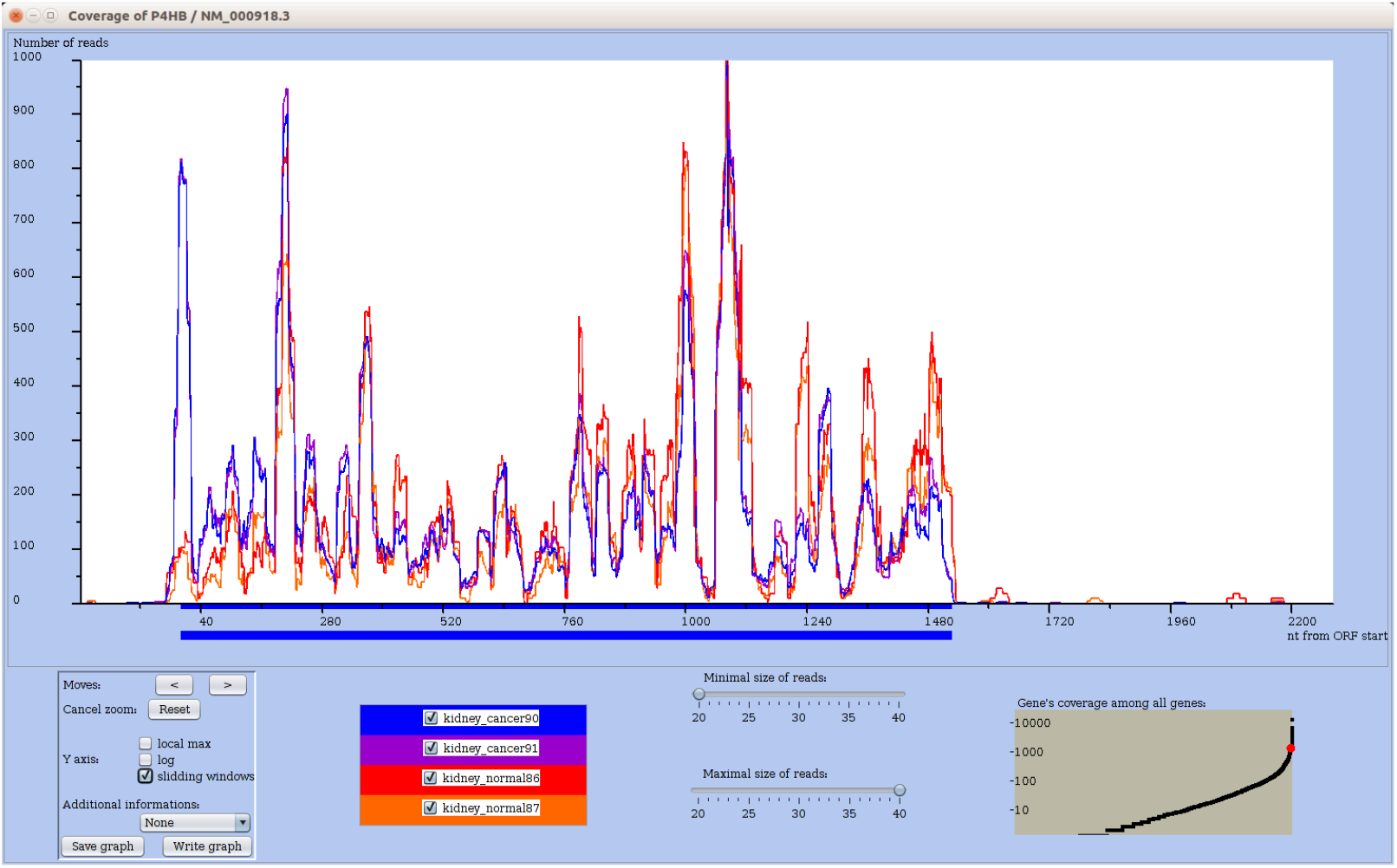
Ribo-seq profiles of a chosen mRNA for multiple conditions. The coding region of the main ORF is materialized by an horizontal blue bar below the profile along the X-axis. The user can interactively change which datasets are plotted, as well as the range of considered read sizes. The plot is updated on-the-fly. Further biological information or annotation, such as measures of codon usage bias, can be displayed on the graph, and the entire graph can be saved as an image file (see User’s manual).

Finally, the user can save the graph as an image (in PNG format) or write down the data of the plot to a tabular file for downstream analyses with a spreadsheet or any scripting language (R, Python).

### 3.4 Data mining queries

RRE provides three major types of comparative queries to identify translational changes between conditions.

1. select subsets of mRNAs according to numerical criteria (e.g. coverage ratio between UTR and CDS regions, RPKM ratio on two conditions, minimal read coverage),
2. detect changes in profile comparison at mRNA level for all mRNAs,
3. find mRNAs with coverage changes in user-defined sub-regions.

Let us illustrate each type by giving one example of biological query.

- Select mRNAs with differential profile within UTRs across conditions (Type 1 query) Here, the user wants to automatically select mRNAs whose main ORF is translated in a given condition, while their UTRs are more covered in the other conditions. Within a dedicated window (Figure 4), the user enters the desired conditions that are combined with logical operations, meanwhile RRE determines the selected subset on the fly (see the User’s manual for a view of the query interface). In our case, the query combines two conditions: i/ a high coverage in the ORF region across all normal datasets (200 reads / along ORF), ii/ a medium coverage on the 5’UTR region in the cancer samples (100 reads in 5’ UTR). RRE yields the list of eight selected mRNAs in table and displays the count for each region in each conditions. The table can be exported for further processing with e.g. any statistical analysis software like R or any spreadsheet. Clicking on an mRNA name, the user inspects the profiles in the desired condition for that gene.
- Find mRNAs with distinct profiles in normal vs tumoral conditions (Type 2 query) The user wants to compare for each mRNA its change in profile across two conditions. In the kidney datasets, the query contrasts a normal vs tumoral conditions. For each mRNA it computes using sliding windows a vector that summarizes the mRNA profile for each condition, and calculates the vector correlation. To discard mRNAs having a coverage too low to get a meaningful correlation, the user can subselect genes with a minimal coverage. RRE displays in the correlation plot, the correlation of each mRNA (on y-axis) by a dot in function of its mean of its total coverage in both conditions). Each dot is clickable to provide the detail on the corresponding mRNA and whose nouns can be added on demand. In Figure 5, most mRNAs get a high correlation, meaning profiles that change mostly in the intensity of the peaks. However, some cases like ATP1B1 exhibits a low correlation suggesting that their profile should differ between the two conditions. Figure 6 shows ATP1B1 profile with 5’UTR and CDS start (up to ~180 bp) regions almost free from reads in normal condition, but highly covered in tumoral condition. Note that all selected genes exhibit changes in profile across conditions, but not necessarily the same changes.
- Detect mRNAs with extended potential translation in their 3’ UTR only under stress (Type 3 query). In a type 1 query, the user can select genes by setting conditions on the profile of annotated regions (UTR, CDS, ORF). However, because annotations may be uncertain or valid in some cell types only, RRE allows a more flexible inspection of user-defined subregions. The user can precisely define the two subregions to compare. In this example, the two regions are the last 200 nucleotides of the CDS and the the first 200 nucleotides of the 3’UTR. For each mRNA, RRE computes the ratio of the read counts for those two regions in each condition. In the plot, each axis measures the ratio in one condition, and each mRNA is represented by a dot positioned according to its ratios in both conditions (Figure 7). Note that here, the plot displays only a subset of all mRNAs: this is made possible by restricting the query on a previously made list of genes. Notice that the “Genes” subpanel of the window has a selection menu to choose among available lists of the session, and for this example the list named “min250inORF” has been selected.

**Figure 4:**
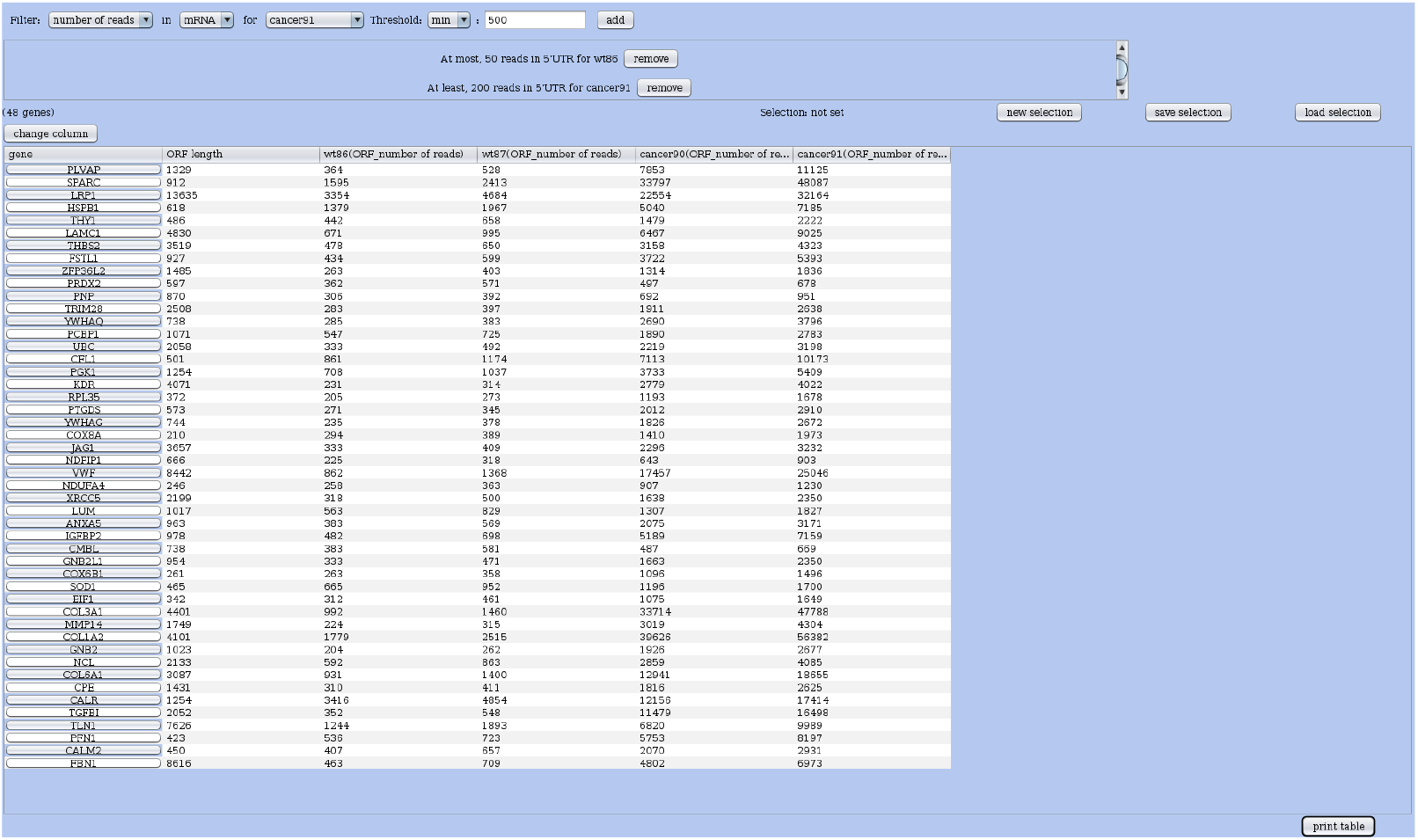
Example of Type 1 query: selecting genes with differential translation between condintions in their 5’UTR. Top: the interactive query editor: the user adds one condition at a time. Below: the table of genes satisfying the query conditions: one gene per line; distinct columns provide the main ORF length, and the read coverages in distinct samples and regions.

**Figure 5:**
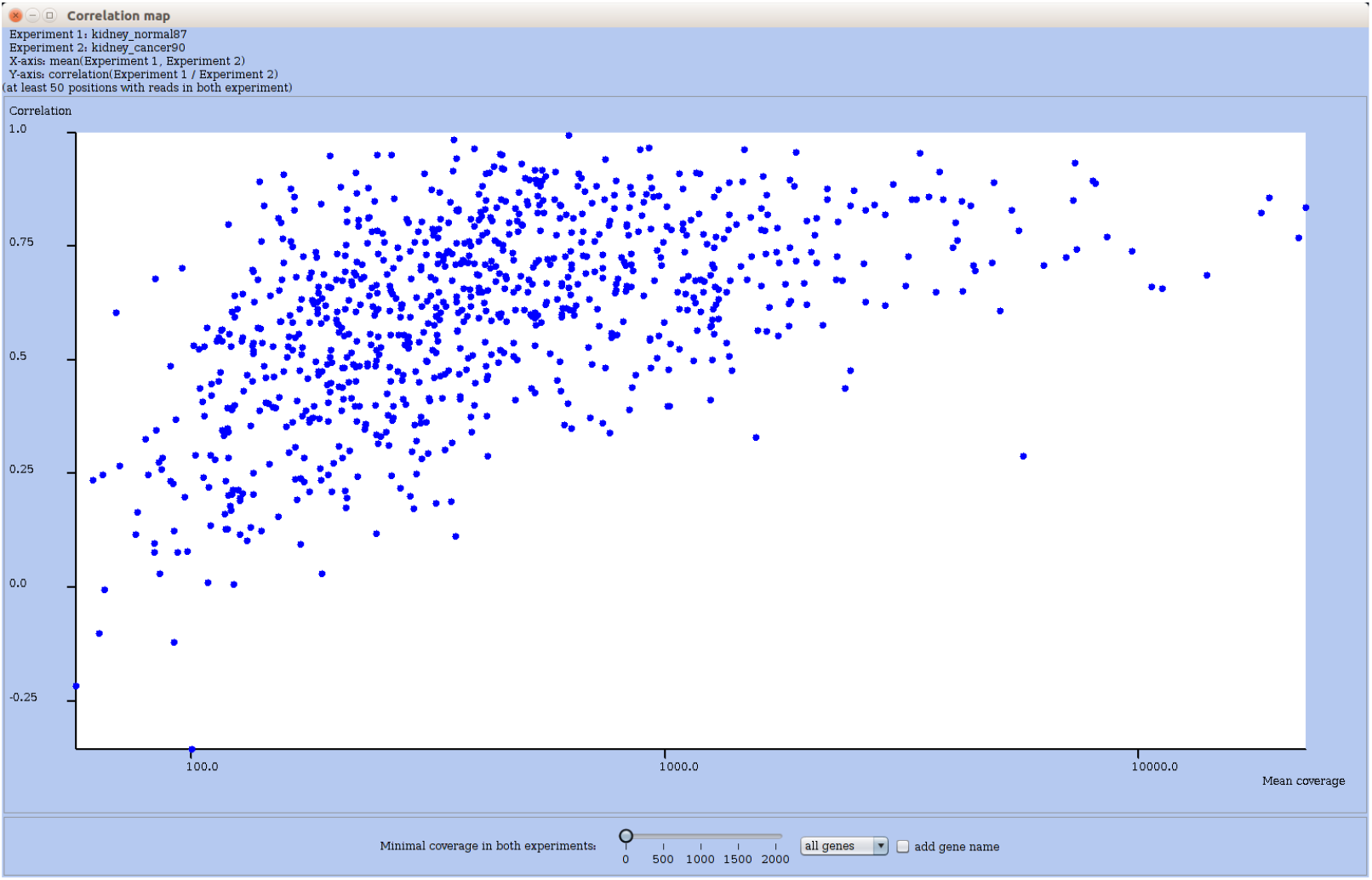
Correlation plot of Ribo-seq profiles of all mRNAs when comparing normal vs tumoral kidney cells. RE plots for each mRNA the correlation of coverages in both conditions (Y-axis) in function of the mean coverage. Each RNA is represented by a dot, and with a click on that dot, one can get the profile coverage for that mRNA in another window. To ease inspection, the user can further refine the set of plotted RNAs by adjusting the coverage threshold: the plot is then updated immediately.

**Figure 6:**
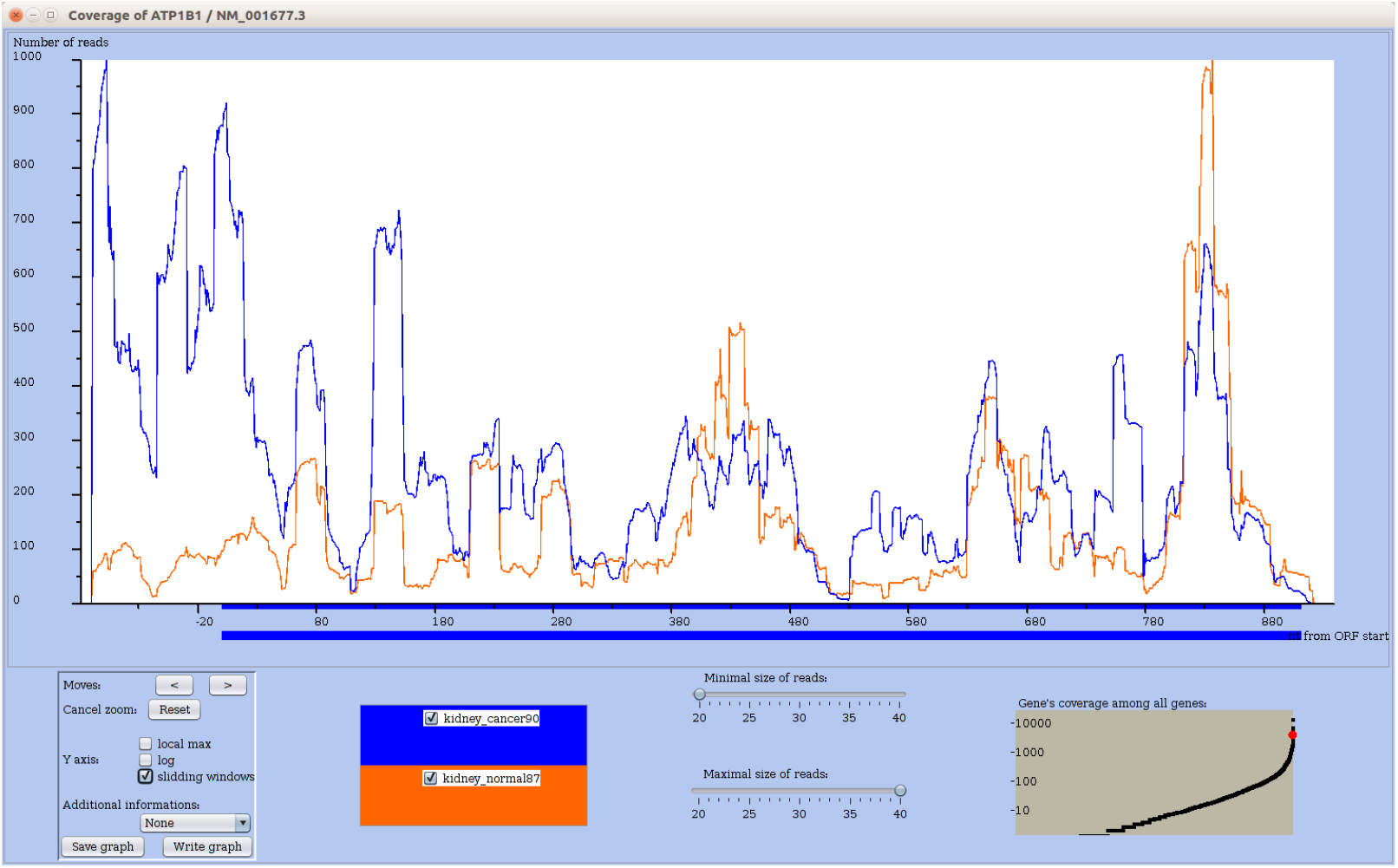
Profile of mRNA ATP1B1, which corresponds to a dot with low correlation between conditions in Figure 5. Sliding window option is on for a better view of the peaks. The normal and cancer conditions differ by the coverage in the 5’UTR of ATP1B1.

**Figure 7:**
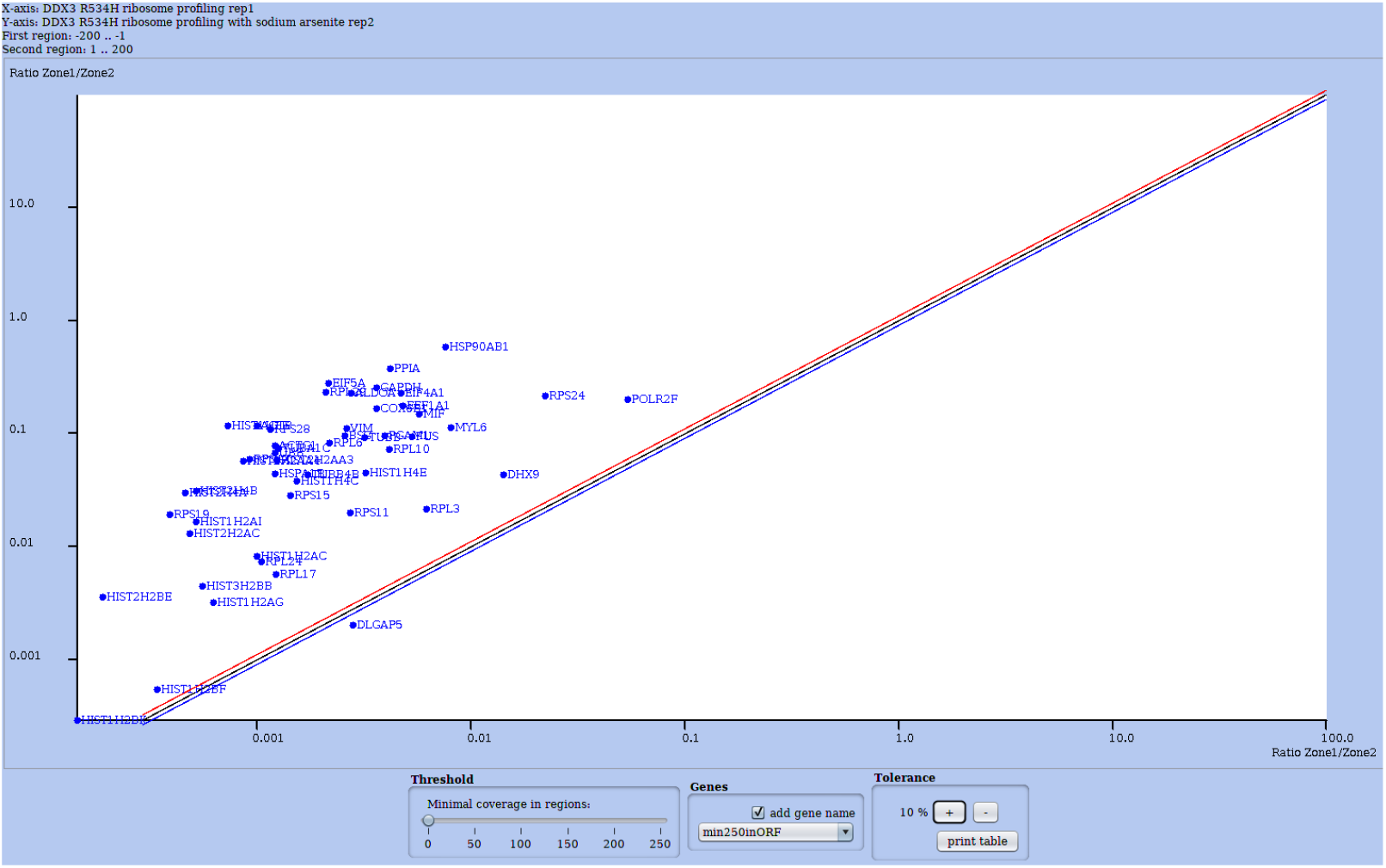
Results of a type 3 query on the HEK dataset: Mining mRNAs with a change in coverage between two user-defined subregions upon treatment with Arsenite. The two regions are the end of CDS and the start of the 3’ UTR (200 nuc. each). The plot is further restricted to a subset of mRNAs (a selection named “min250inORF” in “Genes” subpanel) which was previously obtained by the user with a type 1 query. A dot representing one gene/mRNA indicates its ratio of coverage between the two subregions, in each condition (X-and Y-axis). An example of profile for one mRNA is shown in Figure 8. Like in Figure 5 and in all comparative plots, each RNA is represented by dot and clicking on the dot one can get the profile coverage for that gene. The user can further refine the selection by adjusting the coverage threshold. The main diagonal is bounded by two other diagonals showing a small deviation from a ratio of 1.

In Figure 7, a majority of dots lie above the diagonal meaning that the ratio has increased under stress: for those mRNAs the relative coverage in the two regions is higher when the cells are treated with arsenite. Viewing the profile plot for COX6B1 (Figure 8) shows an example with large peaks within the 3’ UTR only in the arsenite condition. All mRNAs whose dots are above the diagonal (in Figure 7) exhibit an increase coverage in the 200 nuc. of their 3’UTR in the arsenite condition, pointing to an alteration in translation. This illustrates how RRE can help one extracting mRNAs of interest automatically. Note that a gene selected with a type 3 query could be missed with a query of type 1, since the UTR may be large and an alternative ORF may cover only a small portion of it.

**Figure 8:**
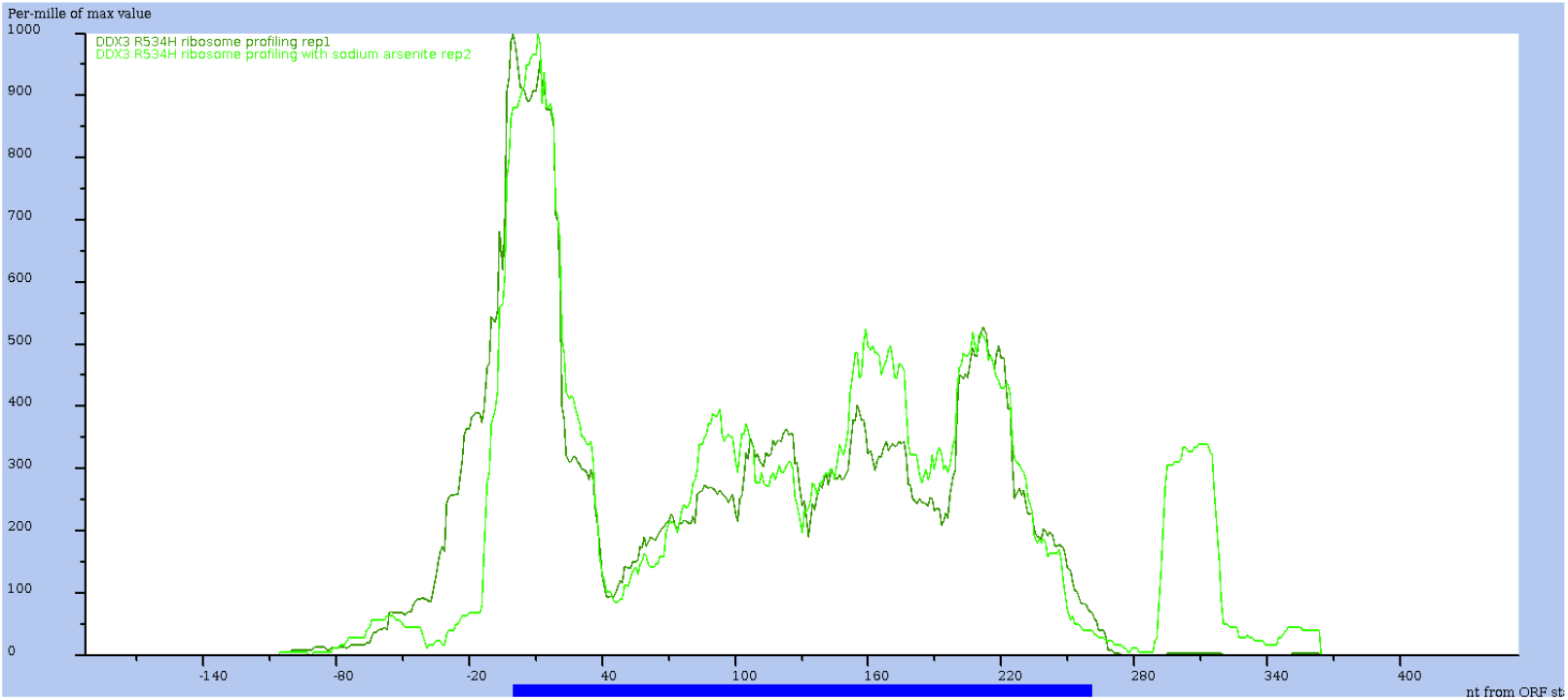
Profile view of mRNA COX6B1, one of the mRNA selected by the type 3 query of Figure 7. Sliding window option is on and the y-axis is relative to the max value in each condition: so the height of the two curves are not comparable (here). One clearly observes a major difference in the 3’UTR: condition with sodium arsenite exhibits a strong relative coverage, while the control condition has almost no reads.

### 3.5 Additional features of RRE

Contrary to other tools, RRE enables the user to restrict on-the-fly the data to a range of read sizes, or to a range of global read coverages, at many steps of the analysis. Using queries the user can save a selection, that is a subset of RNAs of interest, and give it a name. With this selection, general plots illustrated in above Figures or in the manual, can be “restricted” to that subset of RNA interactively. The user can explore its data and save the entire session, which can then be reloaded and continued later. Last, once an analysis has been performed, the user can export graphic figures in PNG format. A comprehensive manual guides the user in all theses manipulations with RRE.

## 4 Conclusion

Despite the publication of numerous methods, programs or R packages for analyzing Ribo-seq [2, 21], a userfriendly, interactive exploration of multiple ribosome profiling dataset remained tedious. Here, we have implemented, made available, and described a standalone tool named RNA-Ribo Explorer (RRE) to offer a solution for mining and visualizing such data to the end user, since Ribo-seq is increasingly used to address a wide variety of questions regarding translational mechanisms and regulation. Our goal was not to re-implement useful R packages or software that perform either read preprocessing (like cleaning and mapping), post-processing or downstream analyses (like differential expression or Ribosomal stalling events), but to ease the visual inspection of the data – which is important for many practitioners. Instead, we opt for a solution that can read in files containing read counts on a reference genome (which are obtained from a preprocessing step) and output graphics or subset of data for downstream more specialized analyses. Hence, RRE can be utilized in intermediate steps of the analysis where interactive exploration is needed. Given the absence of standard ways of analyzing such data (which vary according to the protocol), RRE may well prove a timely and suitable solution. The contribution of RRE is to enable the end user to inspect, compare, and query multiple datasets (either RNA-seq or Ribo-seq) in an easy and flexible manner, and allow adaptable visualization.

Research on translation and epigenetic of RNA (a.k.a. the epitranscriptome) is very active, and require further computational analysis tools than those currently covered by RRE. Beyond read coverage and coding sequence annotation, it would be important to also query sequencing data providing the position of epigenetic marks (e.g., MERIP-seq), known or putative alternative ORFs, or the presence of structural / functional RNA elements (like Internal Ribosomal Entry Sites (IRES) or AU-rich elements (ARE)), just to mention a few complementary data. Such data can already be obtained by sequencing, retrieved from existing databases or from the literature, and analysed on their own, but not in an integrated manner, not through a unified query system like the one provided in RRE. Easy integration of data and flexible querying are key to the end users.

For instance, complex structural RNA elements could be computed or read from annotations, and either added on profile plots or considered in query conditions. This would allow to check whether regions of altered translation (slowdown, stalling) spatially correspond to locations of such structures. Similarly, annotations of potential upstream ORFs could be obtained from a dedicated tool, read from a file, and manipulated within RRE framework. Scores measuring the coding content of a region could be used to predict putative ORFs in mRNAs; in the future, existing measures will be thoroughly evaluated (as suggested in the review [2]), which would ease the choice for incorporating a score measure in RRE.

Finally, information coming from quantitative proteomics or peptidomics would help prioritizing candidate short or alternative ORFs that exhibit coverage in Ribo-seq plots (an example of confirmation of putative peptide encoding ORFs was published in [17]).

## Availability

RRE and its companion tool (FSTC) are both freely available under Cecill license. Both executable software (jar file) and the user’s manual can be downloaded fromhttps://gite.lirmm.fr/rivals/RRE/-/releases. The source code of RRE is available on a gitlab repository at https://gite.lirmm.fr/rivals/RRE. Information and an html version of the user’s manual can be found at http://atgc-montpellier.fr/rre.

## Acknowledgements

This work was supported by the GEM Flagship project funded from Labex NUMEV (ANR-10-LABX-0020), the Institut de Biologie Computationnelle ANR (ANR-11-BINF-0002), by project FluoRib (INCa grant n°2018-131) within the PLBIO program. We thank the ATGC platform for hosting RRE and the Institut Français de Bioinformatique for general support to the platform. Thanks to J. Ripoll for help, to A. Bastide, A. Choquet, and the colleagues in GEM project for feedback on the tool interface.

## Contributions

DP: Conceptualization, Methodology, Software

AD: Funding acquisition, Project administration, Supervision

ER: Conceptualization, Software, Funding acquisition, Project administration, Supervision, Writing

